# Different evolutionary trends form the twilight zone of the bacterial pan-genome

**DOI:** 10.1101/2021.02.15.431222

**Authors:** Gal Horesh, Alyce Taylor-Brown, Stephanie McGimpsey, Florent Lassalle, Jukka Corander, Eva Heinz, Nicholas R. Thomson

## Abstract

The pan-genome is defined as the combined set of all genes in the gene pool of a species. Pan-genome analyses have been very useful in helping to understand different evolutionary dynamics of bacterial species: an open pan-genome often indicates a free-living lifestyle with metabolic versatility, while closed pan-genomes are linked to host-restricted, ecologically specialised bacteria. A detailed understanding of the species pan-genome has also been instrumental in tracking the phylodynamics of emerging drug resistance mechanisms and drug resistant pathogens. However, current approaches to analyse a species’ pan-genome do not take the species population structure into account, nor do they account for the uneven sampling of different lineages, as is commonplace due to over-sampling of clinically relevant representatives. Here we present the application of a population structure-aware approach for classifying genes in a pan-genome based on within-species distribution. We demonstrate our approach on a collection of 7,500 *E. coli* genomes, one of the most-studied bacterial species used as a model for an open pan-genome. We reveal clearly distinct groups of genes, clustered by different underlying evolutionary dynamics, and provide a more biologically informed and accurate description of the species’ pan-genome.

## Main

Advances in whole genome sequencing in the last two decades and the ability to sequence multiple isolates of the same species have revealed that, often, only a small fraction of genes are shared by all species members. Conversely, a substantial proportion of the combined pool of genes within a species – the pan-genome – consists of highly mobile genetic material with heterogeneous distributions across its members (Brockhurst et al. 2019).

In a traditional pan-genome analysis, genes are divided into core genes, describing those present across the majority of the members of the species, and accessory genes, which are only present in some. The accessory genome is often further subdivided into rare and intermediate genes based on their frequency in the dataset. However, measuring gene frequencies across the whole dataset does not account for the population structure or biased sampling of the genomes in the dataset. Such simple classification can be particularly problematic when the population of interest consists of multiple deep-branching lineages that are unevenly represented in the collection. For example, if 50% of a genome collection is represented by one lineage that was heavily over-sampled compared to other lineages, and all isolates of that lineage have a particular gene which is absent in all other lineages, this gene will simply be defined as an “intermediate” gene. Based on these definitions alone, it would not be differentiated from a gene that is found in all isolates of all the other lineages, or evenly distributed across the different lineages comprising 50% of the total isolates. Notably, ecological adaptation of a globally disseminated lineage may be driven by a large set of genes found in all isolates of that lineage, which are rare outside the lineage (Lassalle et al. 2017). Hence, the biological reality requires more refined concepts when classifying genes in the pan-genomic context.

Here, we introduce a population structure-aware approach to classify the genes of a pan-genome beyond accessory and core categories, which accounts for the relative representation of the lineages in the population being studied. This refined classification allows us to better describe the pan-genome and its underlying evolutionary dynamics in organisms with complex population structures. Recent hypotheses on the evolution of the pan-genome have highlighted that different evolutionary mechanisms are required to explain the observed patterns of large open pan-genomes (Vos and Eyre-Walker 2017; Andreani, Hesse, and Vos 2017; Shapiro 2017; McInerney, McNally, and O’Connell 2017). Several competing and non-exclusive hypotheses have been proposed, including the selectively neutral spread of accessory genes – including, but not limited to highly mobile selfish elements (Andreani, Hesse, and Vos 2017; Vos and Eyre-Walker 2017), or indeed adaptive evolution (McInerney, McNally, and O’Connell 2017). Here we illustrate how an analysis of the patterns of within-species gene distribution informed by population structure can provide a more precise view of genes following different evolutionary trajectories. We demonstrate this on a compiled dataset of over 7,500 carefully curated *Escherichia coli* genomes: one of the most-studied bacterial species and used frequently as a model to illustrate an open pan-genome (Touchon et al. 2009; Rasko et al. 2008; Gordienko, Kazanov, and Gelfand 2013).

## Results

### Case study: population structure-aware pan-genome analysis of a collection of 7,500 *E. coli* genomes

To demonstrate how one can refine a pan-genome description while accounting for population structure, we used a recently published genome collection that includes over 7,500 *E. coli* and *Shigella* sp. genomes isolated from human hosts, referred to as the Horesh collection (Horesh et al. 2021). Shigellae are in fact specialised pathotypes of *E. coli* and were thus included (Pettengill, Pettengill, and Binet 2015; Chattaway et al. 2017). Briefly, the genomes in the Horesh collection were collated from publications and other public resources, representing the known diversity of the clinical *E. coli* isolate genomes available in public databases and underwent quality-control steps to ensure a final set of high-quality genomes. The genomes were grouped into lineages of closely related isolates (Figure 1A) using a whole genome-based clustering method that was designed to determine bacterial within-species population structure (Lees et al., 2019.). In total, the collection featured 1,158 lineages representing the *E. coli* species (as described in (Horesh et al. 2021)). We restricted our population-structure aware pan-genome analysis to the largest 47 lineages, which represented the majority of this dataset (7,692/10,158 genomes). Importantly regarding the demonstration of our approach, 70% (5,349/7692) of all genomes in this collection belong to six highly overrepresented lineages. The pan-genome of the Horesh collection was classified into 50,039 homologous gene clusters (as described in (Horesh et al. 2021)).

**Figure 1:**
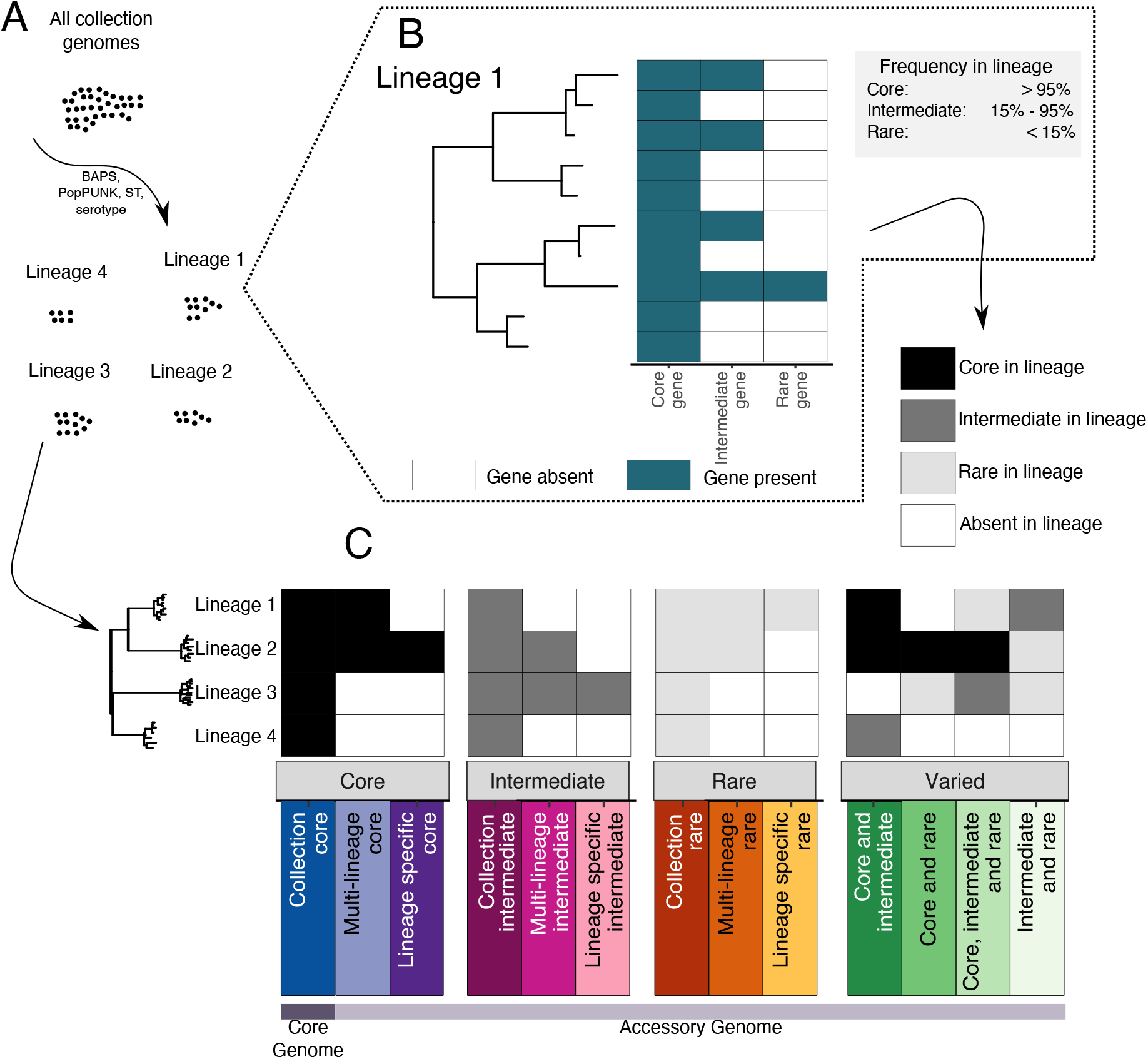
Twilight pan-genome analysis workflow. **A** A collection of genomes are grouped into lineages of closely related isolates. **B** Each gene is classified as core, intermediate or rare in each lineage, depending on its frequency within the lineage (as defined in the grey box). **C** The classification of the entire gene pool across all lineages consists of a total of 13 distribution classes. These include the number of lineages is which a gene is present (all lineages, multiple lineages or a single lineage), and the combination of frequency assignments of the gene in those lineages (core, intermediate or rare).

### The classical definition of the core genome is heavily influenced by the underlying biases of the studied datasets

We defined the distribution for each gene cluster in the *E. coli* and *Shigella* genome dataset by considering their frequency in each of the above-defined lineages independently. A gene cluster can thus be core, intermediate, rare or absent based on its frequency within each respective lineage (Figure 1B) but can have varied distributions in different lineages (Figure 1C, e.g. core in some and rare in other lineages). We summarised the combination of gene cluster occurrence patterns across lineages into a set of 13 species-wide distribution patterns, which we propose as novel categories for a more appropriate description of datasets with complex underlying population structure (Figure 1C). Compared to traditional pan-genome analyses, the “collection core” genes represent the classical definition of the core genome, whereas we consider the accessory genome as subdivided into 12 new classes, informed by the population structure, whose distribution reflects several different evolutionary dynamics.

Figure 2A illustrates the new distribution classes, based on the number of lineages in which they were observed and their mean frequency within those lineages. Only the top right corner represents the traditional set of core genes. The rest of the pane is what is usually summarised as the accessory genome; the colours describe the underlying distribution classes. The plot shows the continuity of gene frequencies across the entire collection, with genes present across almost the entire distribution frequency spectrum.

**Figure 2:**
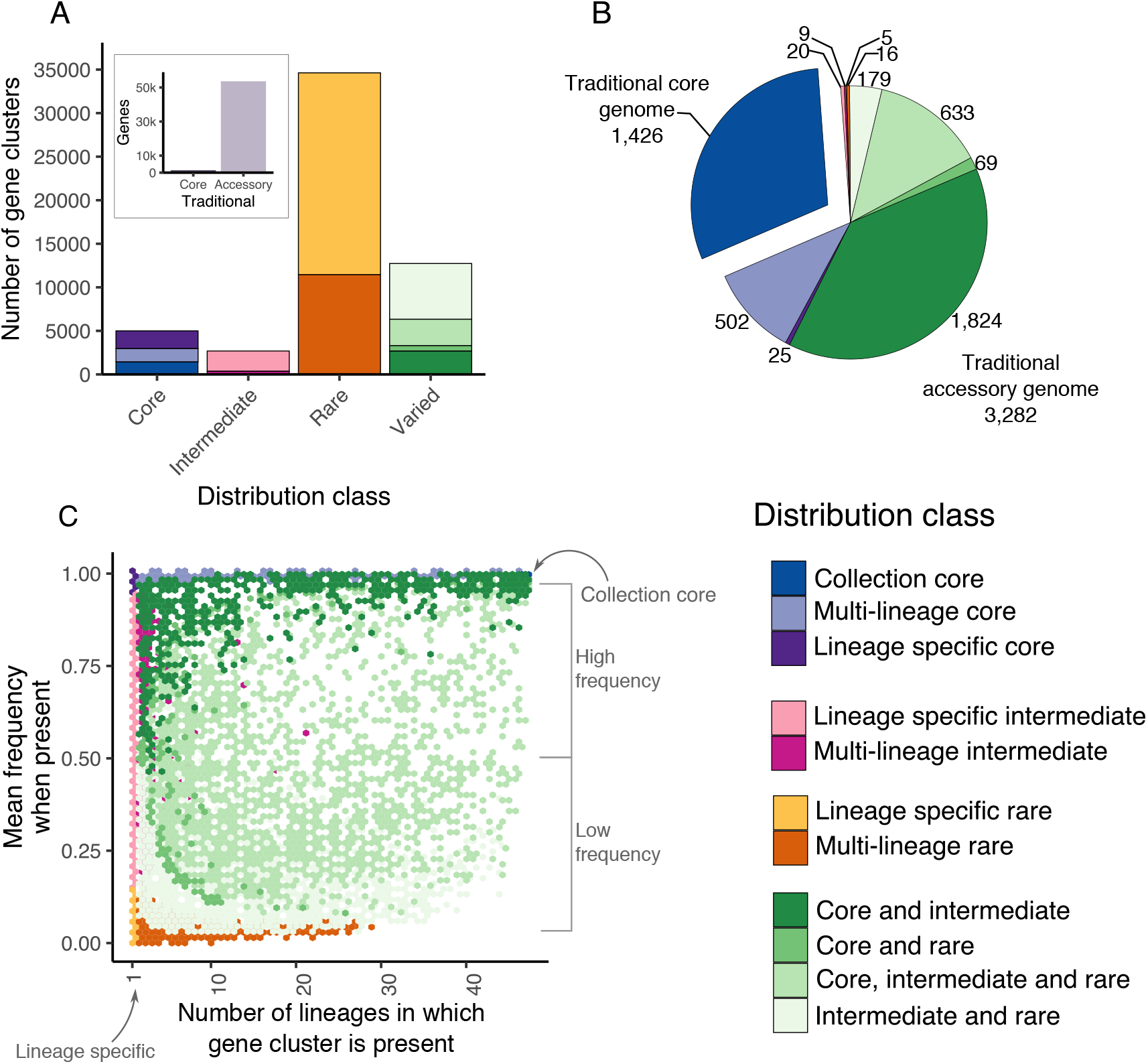
Population-structure aware pan-genome of E. coli. **A** Hexagonal binning of all genes of the E. coli pan-genome, presented as the number of lineages in which each gene was observed (x-axis) against the mean frequency across the lineages containing it (y-axis). Each hexagon is coloured by the most common distribution class on the pane (see colour key). **B** Number of gene clusters of the E. coli pan-genome from each of the novel distribution classes. **C** The relative abundance and gene count of each of the distribution classes in a typical E. coli genome in the collection. Only the collection core genes represent the traditional set of core genes, the rest represent what would usually all be summarised as the accessory genome.

Within this expanded classification, “collection core genes” are equivalent to the traditional classification of core (assuming a threshold of ≥95% of the genomes in the collection encoding for a gene for it to be defined as core). In this analysis, the collection core is comprised of 1,426 gene clusters; representing 3% of the total number of gene clusters comprising the *E. coli* pan-genome (1,426/50,039) and 30% of the total number of genes in a typical *E. coli* genome (defined as the weighted median across the 47 lineages, see methods, Figure 2B,C, Supplementary Table S1).

An additional 1,532 gene clusters (3% of the pan-genome) are now defined as multi-lineage core: that is, they are present in ≥95% of isolates per lineage in multiple (but not all) lineages (2-46 lineages, Figure 2B). Another 2,040 genes (4% of all genes) were core to only a single lineage (Figure 2B). Both classes would have been assigned to the accessory genome following the classical definition of the pan-genome, as genes that are core to lineages with low representation in the dataset would have been categorised as rare genes. Importantly, these two additional distribution classes allow us to capture more recent acquisition or loss events that have remained fixed in a respective lineage or lineages.

### The majority of rare and intermediate genes are lineage-specific

The majority of the *E. coli* gene clusters were classified as “rare genes” (Figure 2B, defined as present in <15% of isolates of a lineage) in one or multiple lineages within the dataset. In total, 63% (34,624/55,039) of the *E. coli* pan-genome was classified as rare, with 67% of all rare genes being specific to a single lineage (23,175/34,624; Figure 2B). In relation to a single *E. coli* genome, these genes only form 0.1% of a typical genome (Figure 2C).

Intermediate frequency gene clusters on the contrary formed only 4% (2,685/55,039) of the entire gene pool; however, similar to the rare gene clusters, 86% of intermediate gene clusters (2,329/2,685) were only observed in a single lineage. Rare and intermediate genes observed in multiple lineages were most commonly observed in up to four lineages (Figure 2C, Supplementary Figure S1). We did not observe any rare or intermediate genes present across more than 30 lineages, and there were no collection rare or collection intermediate genes in this dataset (Figure 1A, 2A,B, Supplementary Figure S1).

### A fifth of the pan-genome consists of genes observed in different frequencies across the lineages

“Varied genes” were defined as those observed in several lineages, but at different frequencies within the respective lineages (e.g. core in one and intermediate in another lineage). These represented 23% of the pan-genome (12,732/55,039) (Figure 2B) and 57% of all genes in a typical *E. coli* genome (Figure 2C). To summarise all of these observations, genes were categorised as “core and intermediate”, “core, intermediate and rare”, “core and rare” or “intermediate and rare” depending on the combination of frequencies in which they appeared (Figure 1C). “Core and intermediate” genes were commonly observed in more lineages and in higher frequencies within those lineages and represented 38% of the genes in a typical *E. coli* genome (Figure 2A,C Supplementary Figure S1). On the other hand, the group of “intermediate and rare’’ had a lower frequency and were observed in fewer lineages (Figure 2A, Supplementary Figure S1).

### Low frequency genes are four times more likely to have been horizontally transferred than high frequency genes

As the pan-genome in any collection represents a snapshot of the gene pool at the time of sampling, our refined view of the different distribution classes may be used to infer how the genes are gained and lost and can indicate a gene’s future trajectory within a population. For instance, genes that are self-mobile or carried as cargo on mobile genetic elements will have a markedly different pattern of distribution relative to genes that may be in the process of being selectively lost in any particular lineage.

To assess whether genes from the different distribution classes showed varying evidence of levels of mobility and estimate the probability of genes having been horizontally transferred, we applied a species-tree gene-tree reconciliation method (Morel et al. 2020) to each gene cluster of the pan-genome. As expected, higher frequency genes (Figure 2B), ie. those present in the “collection core”, “core and intermediate” and “multi-lineage core”, gene sets were estimated to have the lowest probabilities of having been horizontally transferred (median 0.12, 0.13 and 0.1, respectively) (Figure 3A, Supplementary Figure S2). Conversely, the lower frequency gene classes, i.e. “multi-lineage rare”, “multi-lineage intermediate”, “intermediate and rare” and “core, intermediate and rare” gene sets were estimated to be up to four times more likely to have been horizontally transferred than the high frequency genes (median probabilities of 0.48, 0.46, 0.44 and 0.31, respectively, Supplementary Figure S2). Consistent with this, by counting the total number of gene gain events predicted to have occurred on each branch using ancestral state-reconstruction, multi-lineage core gene gains most commonly occurred along the internal branches (Figure 3B) whereas “intermediate and rare” genes were predominantly gained at the branch tips (Figure 3C).

**Figure 3:**
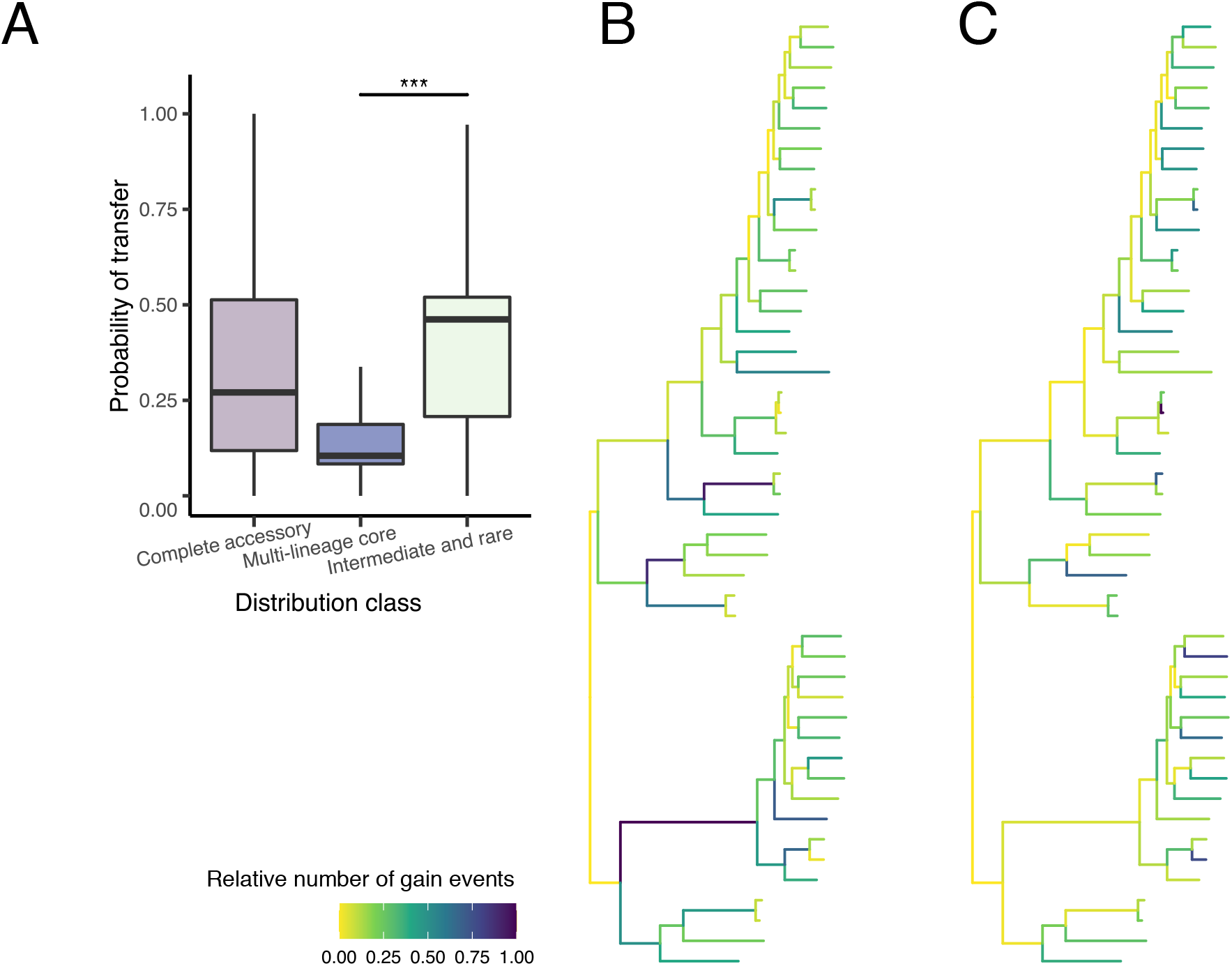
Different evolutionary dynamics of genes within the accessory genome. **A** Inferred probability of transfer using species-tree gene-tree reconciliation for the entire accessory genome (i.e. all 12 distribution classes which make up the accessory genome), only the “multi-lineage core” genes, and only ‘intermediate and rare’ genes (Wilcoxon rank sum test, ***p < 0.001). **B,C** number of gain events estimated to have occurred on each branch using ancestral state reconstruction when considering the ‘multi-lineage core’ genes (**B**) or all the ‘intermediate and rare’ genes (**C**). Darker colours represent more gain events were estimated to have occurred on a branch.

Of the multi-lineage core genes, 54% could be assigned as basic cellular processes such as metabolism, information storage and processing and cell signalling (Supplementary Figure S3). On the other hand, 73% of “intermediate and rare” genes were either assigned to a poorly characterised function (often associated with genetic mobility) or of unknown function (Supplementary Figure S4).

### Detection of shared horizontally transferred genes between lineages is strongly dependent on unbiased sampling

We observed that the number of “intermediate and rare” genes shared between every two lineages was positively correlated with the size of the two lineages being compared, with larger lineages sharing more mobile genes (Figure 4A, log linear regression, *R^2^*=0.45, *p*<2.2e-16). Contrarily, we did not observe a relationship between the number of “intermediate and rare” genes shared between every two lineages and their phylogenetic distance (Figure 4B; linear regression, *R^2^*=0.005, *p*=0.01). Using our population-structure aware approach to measure sharing of the genes belonging to the different distribution classes suggests a lack of barrier to gene flow between lineages. With that being said, our analysis highlights the need to increase sampling of under-studied lineages in order to overcome sampling-related biases and truly understand the level of horizontal transfer of genes between them.

**Figure 4:**
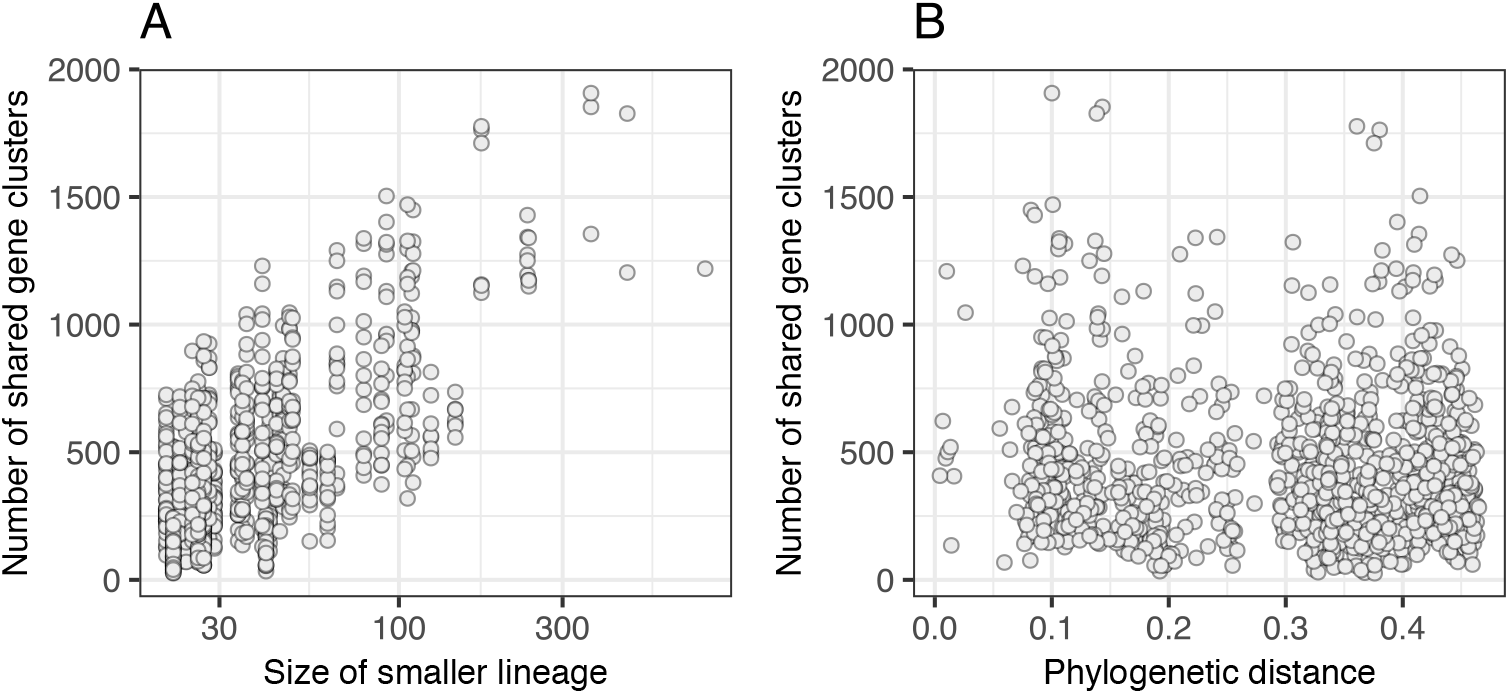
Relationship between sharing of “intermediate and rare” genes, phylogenetic distance and lineage size. Relationship between the number of “intermediate and rare” genes shared between every two lineages and the size of the smaller lineage of the two being compared (**A**) or the phylogenetic distance between them (**B**). Pairwise comparisons were considered between every two of the 47 lineages.

### Novel distribution classes can highlight lineages with evolutionary trajectories unusual for the species

We normalised the counts of shared genes to correct for the bias led by the size of the lineages and any sharing of genes driven by phylogenetic relatedness (see Methods, Supplementary Figure S5). This revealed that two lineages (12 and 40) tended to share more “intermediate and rare” genes than expected compared to other lineages in the collection (Pairwise Wilcoxon rank sum test, p<0.001, FDR corrected, Figure 5A, Supplementary Figure S6). Genomes in lineages 12 and 40 however, are smaller than those in other lineages (Pairwise Wilcoxon rank sum test, p<0.001, FDR corrected, Figure 5B), and the mean number of lineage-specific rare genes in a single genome was 32 and 30 genes, respectively, compared to 5 in a typical *E. coli* genome (Pairwise Wilcoxon rank sum test, p<0.001, FDR corrected; Figure 2C, Figure 5C, Supplementary Figure S7). Overall, the relative fraction of lineage-specific rare genes in the genomes of these lineages was seven times higher relative to the median fraction in the entire collection (median fraction in collection = 0.001; median fraction in lineages 12 and 40: 0.007; Figure 2C). Similar to the other low frequency genes, the “lineage-specific rare” genes were also most commonly predicted to be phage-derived or otherwise had other annotations related to genetic mobility (Supplementary Figure S4).

**Figure 5:**
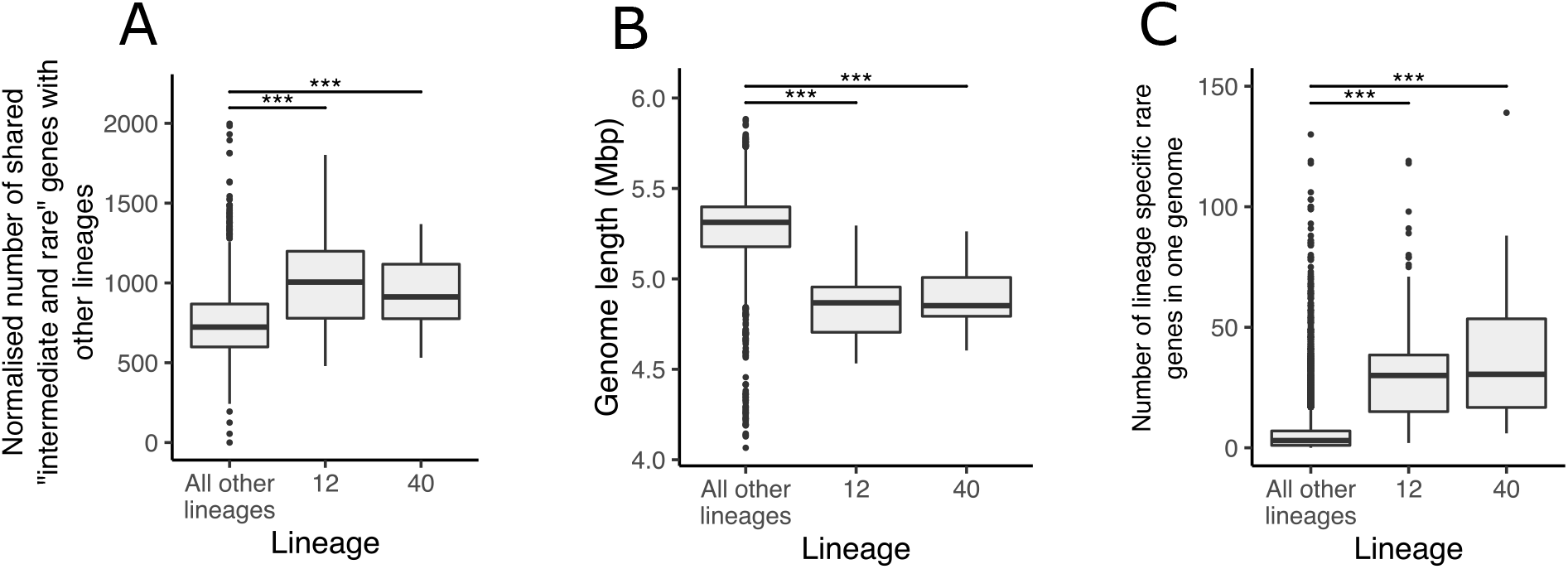
Redefining the pan-genome reveals key insights into particular lineages. **A** Number of shared mobile genes per isolate, for isolates belonging to lineage 12, 40 or all other lineages. Counts were normalised to consider the dependency on the lineage size, and to correct for gene sharing driven by phylogenetic relatedness (Pairwise Wilcoxon rank sum test, FDR corrected, *p < 0.05, **p< 0.01, ***p < 0.001). **B** Genome length of each isolate, for isolates belonging to lineage 12, 40 and all other lineages (Pairwise Wilcoxon rank sum test, FDR corrected, *p < 0.05, **p< 0.01, ***p < 0.001). **C** Number of “lineage specific rare” genes observed in each isolate, for isolates belonging to lineage 12, 40 and all other lineages. (Pairwise Wilcoxon rank sum test, FDR corrected, *p < 0.05, **p< 0.01, ***p < 0.001).

## Discussion

To date, the existence of complex population structure and diverse lineages in the bacterial populations has not been taken into account in pan-genome analyses. We introduce a population-structure aware classification of the pan-genome as an extended set of thirteen classes. Our study reveals distinctive patterns in the evolutionary dynamics of these gene classes, with differences in the relative importance of these gene classes between lineages within *E. coli*. Our approach can be further applied to other bacterial species of public health interest to provide insight into the evolutionary dynamics of genes within such species.

Subcategorising the genes of the accessory genome allowed us to distinguish the evolutionary dynamics of different gene classes within the accessory genome. Grouping all the genes of the accessory genome together showed a large spread of probabilities of genes being horizontally transferred. Our refined approach showed that low-frequency genes transfer more frequently than the high-frequency genes. Importantly, the study of outliers, which disagree with the general trend of each of the distribution classes, can reveal gene-specific evolutionary dynamics, including adaptive processes. For instance, multi-lineage core genes estimated to have high rates of transfer may represent genes that were acquired and fixed independently on multiple occasions and could be cases of convergent evolution and adaptation to similar niches.

By expanding the number of distribution classes of the accessory genome relative to traditional approaches, we were able to observe a relationship between the number of rare genes per genome and high levels of sharing of horizontally transferred genes in two lineages, 12 and 40. This relationship has biological implications, as it suggests that the higher levels of gene sharing are driven by an increased ability to retain mobile genes in each genome for isolates belonging to these lineages, or an inability to prevent invasion by foreign selfish elements. 78% of the isolates from lineage 12 are of ST10 and 43% of the isolates in lineage 40 are from ST23. ST10 and ST23 are ubiquitous as they have been described as both commensal and pathogenic, multidrug resistant, as well as isolated from human and animal sources (Bortolaia et al. 2011; Oteo et al. 2009). These properties have labelled these lineages as generalists and as potential facilitators of gene movement in the population (Matamoros et al. 2017). Here we showed that these differences can be identified and exemplified through more refined analysis of the pan-genome of the entire dataset, as well as within each lineage separately. In doing so, we can also identify lineages that have a greater propensity as vectors for facilitating gene movement.

It is clear that as available genomic data grows, and our understanding of the population structure becomes richer, a population structure-aware approach to analysing the gene frequency distribution is necessary to overcome several biases inherent in large datasets consisting of variably sampled populations, as these biases can overshadow the true distribution of the genes in a population. For example, using a traditional approach, treating all gene counts across the entire collection equally, genes that are core and specific to a single lineage that has a low representation or penetrance in the collection could be mistaken for rare genes. Identification of these genes is highly important, as being core to only a subset of the population suggests that they have an evolutionary advantage in a particular genetic context or ecological setting (Lassalle, Muller, and Nesme 2015; Gori et al. 2020). Additionally, genes that are core to a subset of the population are particularly relevant to investigate further for their potential use in diagnostics and epidemiology.

## Materials and methods

### Gene classification into “distribution classes”

Each gene cluster was assigned to a distribution class based on its frequency within genomes belonging to the same phylogenetic clusters, termed lineages (Figure 1A). Within each lineage, a gene was defined as “core” if it was present in more than 95% of the isolates of that lineage, “intermediate” if present in 15% to 95% of isolates of the lineage, and “rare” if present in up to 15% of the isolates of the lineage (Figure 1B). Three main distribution classes, “Core”, “Intermediate” and “Rare”, contained all the genes that were always observed as being “core”, “intermediate” or “rare” respectively across the lineages in which they were present (Figure 1C). “Collection core”, “collection intermediate” and “collection rare” genes were present and in their respective frequencies across all the lineages of the collection. “Multi-lineage core”, “multi-lineage intermediate” and “multi-lineage rare” genes were present in multiple lineages in their respective frequencies. “Lineage specific core”, “lineage specific intermediate” and “lineage specific rare” genes were present only in one lineage in their respective frequencies. The final main distribution class or “varied” genes, included all the genes which were observed as either combination of “core”, “intermediate” or “rare” across multiple lineages. All the possible combinations are “core, intermediate and rare”, “core and intermediate”, “core and rare” and “intermediate and rare” (Figure 1C). The classification of all genes in the *E. coli* collection is available as Supplementary Table S1.

### Measuring the genetic composition of each lineage

The number of genes from each of the thirteen distribution classes was counted in each of the 7,693 *E. coli* genomes in the collection. The median number of genes from each distribution class was calculated per lineage. The genetic composition of a typical *E. coli* genome was measured as the median across the medians calculated per lineage for each distribution class.

### Gene-tree species-tree reconciliation

GeneRax (v1.2.2) was used to infer the probability of a horizontal gene transfer event for each gene using species-tree gene-tree reconciliation (Morel et al. 2020). A multiple sequence alignment of all the representative sequences of each gene cluster which had at least four members (available as file F6 at (Horesh et al. 2021)) were performed using mafft (v7.310) (Katoh and Standley 2013). An initial tree for each gene cluster, used as the input for GeneRax, was constructed using iqtree (v1.6.10) with SH-like approximate likelihood ratio test (SH-aLRT) with 1000 replicates (Nguyen et al. 2015). The reconciliation was performed against the species tree provided in (Horesh et al. 2021) with strategy SPR, reconciliation model UndatedDTL and substitution model GTR+G. The probability of transfer was inferred by GeneRax for each of the gene-clusters when reconciled against the species tree.

### Counting gain events

The phylogenetic tree representing the 47 lineages was downloaded from (Horesh et al. 2021). The phylogenetic distance between every two lineages was measured as the patristic distance using the function ‘cophenetic’ from the R package ape (v5.3) (Paradis, Claude, and Strimmer 2004). The patristic distance is the sum of the total distance between two leaves of the tree, which represent the lineages, and hence summarises the total genetic change in the core gene alignment represented in the tree.

The leaves or tips of the phylogenetic tree represent the 47 lineages. Presence of a gene in a lineage (tree leaf) was defined as the gene being observed at least once in at least one isolate of the lineage, i.e. the frequency in the lineage was ignored. The presence or absence of a gene in an ancestral node, i.e. an internal node, was determined using accelerated transformation (ACCTRAN) reconstruction implemented in R (Farris 1970). ACCTRAN is a maximum parsimony-based approach which minimises the number of transition events on the tree (from absence to presence and vice versa) while preferring changes along tree branches closer to the root of the tree.

Gain and loss events were counted based on the results of the ancestral state reconstruction. If there was a change from absence to presence from an ancestor to a child along a branch in the phylogeny, a gain event was counted. If there was a change from presence to absence a loss event was counted. The total number of gain and loss events was counted for each gene as well as on each branch for all distribution classes. ggtree (v1.16.6) was used for phylogenetic visualisation (Yu et al. 2017).

### Measuring gene sharing between lineages

The number of genes shared from each distribution class between every two lineages was counted using custom R and python scripts. In order to identify where some lineages shared more genes than expected, we corrected for gene sharing driven by the phylogeny or by a large sample size. To correct for phylogenetically driven gene sharing, for each lineage we only counted the number of genes shared with lineages which had a patristic distance of 0.15 or more from it on the species tree. This threshold was chosen based on the observation that isolates from the same phylogroup had a patristic distance lower than 0.15 (Supplementary Figure 4). To correct for the lineage size, we fitted a linear model for the number of genes shared between every two lineages against the size of the lineage, which showed a positive coefficient. We adjusted the values as follows: *counts_new_ = count_orig_* – *β × log10(size) – α*, where *β* is the coefficient of the line and *α* is the intercept (Supplementary Figure S5). We then scaled the numbers to be larger than 0 by adding the lowest value to all counts. The new counts no longer correlated with the size of the lineages (Supplementary Figure S5).

### Functional assignment of COG categories

The predicted function and COG category of each gene cluster were assigned using eggNOG-mapper (1.0.3) on the representative sequence of each of the gene clusters (Huerta-Cepas et al. 2017). Diamond was used for a fast local protein alignment of the representative sequences against the eggNOG protein database (implemented within eggNOG-mapper). The COG (Clusters of Orthologous Groups) classification scheme comprises 22 COG categories which are broadly divided into functions relating to cellular processes and signaling, information storage and processing, metabolism and genes which are poorly categorised (Galperin et al. 2015). When no match was found in the eggNOG database, the genes were marked as “?” in their COG category.

Sub-sentences of all lengths were extracted from each of the functional predictions for each gene cluster using the function “combinations” from the python package “itertools”, while ignoring common words. For instance, for the functional prediction “atp-binding component of a transport system”, the words “of”, “a” and “system” were ignored, and the extracted sub-sentences were “atp-binding component”, “atp-binding component transport” and “component transport”. The number of times each sub-sentence appeared in each distribution class was counted. Overlapping sub-sentences which only had a difference of 3 or smaller in their total counts per distribution class were merged in the final count to include only the longer sub-sentence. For instance, if “atp-binding component transport” was counted 100 times and “atp-binding component” was counted 103 times, the final count would only include the longer sub-sentence “atp-binding component transport” with a count of 100.

### Code availability

All analyses were performed using custom R and Python scripts, available at https://github.com/ghoresh11/twilight/tree/master/manuscript_scripts. The script used to classify the genes into distribution classes and generate the figures presented in this study is available at https://github.com/ghoresh11/twilight. The script can be applied on any other dataset, given a gene presence absence file as generated by pan-genome analysis tools and a grouping of each genome into a lineage. ggplot2 was used for all plotting (Wickham 2016).

## Acknowledgements

This work was funded by Wellcome Sanger Institute (no. 206194), a Wellcome Sanger Institute PhD studentship (to G.H. and S.M.), Woolf Fisher Scholarship (to S.M.) and an ERC grant (742158 to J.C). EH acknowledges funding from Wellcome (217303/Z/19/Z) and UKRI (BBSRC V011278/1). We would like to thank Leopold Parts, Simon Harris, Andres Floto and members of the Thomson team for useful discussions.

## References

Andreani, Nadia Andrea, Elze Hesse, and Michiel Vos. 2017. “Prokaryote Genome Fluidity Is Dependent on Effective Population Size.” The ISME Journal 11 (7):1719–21.

Bortolaia, Valeria, Jesper Larsen, Peter Damborg, and Luca Guardabassi. 2011. “Potential Pathogenicity and Host Range of Extended-Spectrum Beta-Lactamase-Producing Escherichia Coli Isolates from Healthy Poultry.” Applied and Environmental Microbiology 77 (16):5830–33.

Brockhurst, Michael A., Ellie Harrison, James P. J. Hall, Thomas Richards, Alan McNally, and Craig MacLean. 2019. “The Ecology and Evolution of Pangenomes.” Current Biology: CB 29 (20):R1094–1103.

Chattaway, Marie A., Ulf Schaefer, Rediat Tewolde, Timothy J. Dallman, and Claire Jenkins. 2017. “Identification of Escherichia Coli and Shigella Species from Whole-Genome Sequences.” Journal of Clinical Microbiology 55 (2):616–23.

Farris, James S. 1970. “Methods for Computing Wagner Trees.” Systematic Biology 19 (1):83–92.

Galperin, Michael Y., Kira S. Makarova, Yuri I. Wolf, and Eugene V. Koonin. 2015. “Expanded Microbial Genome Coverage and Improved Protein Family Annotation in the COG Database.” Nucleic Acids Research 43 (Database issue):D261–69.

Gordienko, Evgeny N., Marat D. Kazanov, and Mikhail S. Gelfand. 2013. “Evolution of Pan-Genomes of Escherichia Coli, Shigella Spp., and Salmonella Enterica.” Journal of Bacteriology 195 (12):2786–92.

Gori, Andrea, Odile B. Harrison, Ethwako Mlia, Yo Nishihara, Jia Mun Chan, Jacquline Msefula, Macpherson Mallewa, et al. 2020. “Pan-GWAS of Streptococcus Agalactiae Highlights Lineage-Specific Genes Associated with Virulence and Niche Adaptation.” mBio 11 (3). https://doi.org/10.1128/mBio.00728-20.

Horesh, Gal, Grace A. Blackwell, Gerry Tonkin-Hill, Jukka Corander, Eva Heinz, and Nicholas R. Thomson. 2021. “A Comprehensive and High-Quality Collection of Escherichia Coli Genomes and Their Genes.” Microbial Genomics, January. https://doi.org/10.1099/mgen.0.000499.

Huerta-Cepas, Jaime, Kristoffer Forslund, Luis Pedro Coelho, Damian Szklarczyk, Lars Juhl Jensen, Christian von Mering, and Peer Bork. 2017. “Fast Genome-Wide Functional Annotation through Orthology Assignment by eggNOG-Mapper.” Molecular Biology and Evolution 34 (8):2115–22.

Katoh, Kazutaka, and Daron M. Standley. 2013. “MAFFT Multiple Sequence Alignment Software Version 7: Improvements in Performance and Usability.” Molecular Biology and Evolution 30 (4):772–80.

Lassalle, Florent, Daniel Muller, and Xavier Nesme. 2015. “Ecological Speciation in Bacteria: Reverse Ecology Approaches Reveal the Adaptive Part of Bacterial Cladogenesis.” Research in Microbiology 166 (10):729–41.

Lassalle, Florent, Rémi Planel, Simon Penel, David Chapulliot, Valérie Barbe, Audrey Dubost, Alexandra Calteau, et al. 2017. “Ancestral Genome Estimation Reveals the History of Ecological Diversification in Agrobacterium.” Genome Biology and Evolution 9 (12):3413–31.

Lees, John A., Simon R. Harris, Gerry Tonkin-Hill, Rebecca A. Gladstone, Stephanie W. Lo, Jeffrey N. Weiser, Jukka Corander, Stephen D. Bentley, and Nicholas J. Croucher. 2019. “Fast and Flexible Bacterial Genomic Epidemiology with PopPUNK.” https://doi.org/10.1101/360917.

Matamoros, Sébastien, Jarne M. van Hattem, Maris S. Arcilla, Niels Willemse, Damian C. Melles, John Penders, Trung Nguyen Vinh, et al. 2017. “Global Phylogenetic Analysis of Escherichia Coli and Plasmids Carrying the Mcr-1 Gene Indicates Bacterial Diversity but Plasmid Restriction.” Scientific Reports 7 (1): 15364.

McInerney, James O., Alan McNally, and Mary J. O’Connell. 2017. “Why Prokaryotes Have Pangenomes.” Nature Microbiology 2 (March): 17040.

Morel, Benoit, Alexey M. Kozlov, Alexandros Stamatakis, and Gergely J. Szöllősi. 2020. “GeneRax: A Tool for Species Tree-Aware Maximum Likelihood Based Gene Family Tree Inference under Gene Duplication, Transfer, and Loss.” Molecular Biology and Evolution, June. https://doi.org/10.1093/molbev/msaa141.

Nguyen, Lam-Tung, Heiko A. Schmidt, Arndt von Haeseler, and Bui Quang Minh. 2015. “IQ-TREE: A Fast and Effective Stochastic Algorithm for Estimating Maximum-Likelihood Phylogenies.” Molecular Biology and Evolution 32 (1):268–74.

Oteo, Jesús, Karol Diestra, Carlos Juan, Verónica Bautista, Angela Novais, María Pérez-Vázquez, Bartolome Moyá, et al. 2009. “Extended-Spectrum Beta-Lactamase-Producing Escherichia Coli in Spain Belong to a Large Variety of Multilocus Sequence Typing Types, Including ST10 complex/A, ST23 complex/A and ST131/B2.” International Journal of Antimicrobial Agents 34 (2):173–76.

Paradis, Emmanuel, Julien Claude, and Korbinian Strimmer. 2004. “APE: Analyses of Phylogenetics and Evolution in R Language.” Bioinformatics 20 (2):289–90.

Pettengill, Emily A., James B. Pettengill, and Rachel Binet. 2015. “Phylogenetic Analyses of Shigella and Enteroinvasive Escherichia Coli for the Identification of Molecular Epidemiological Markers: Whole-Genome Comparative Analysis Does Not Support Distinct Genera Designation.” Frontiers in Microbiology 6: 1573.

Rasko, David A., M. J. Rosovitz, Garry S. A. Myers, Emmanuel F. Mongodin, W. Florian Fricke, Pawel Gajer, Jonathan Crabtree, et al. 2008. “The Pangenome Structure of Escherichia Coli: Comparative Genomic Analysis of E. Coli Commensal and Pathogenic Isolates.” Journal of Bacteriology 190 (20):6881–93.

Shapiro, B. Jesse. 2017. “The Population Genetics of Pangenomes.” Nature Microbiology.

Touchon, Marie, Claire Hoede, Olivier Tenaillon, Valérie Barbe, Simon Baeriswyl, Philippe Bidet, Edouard Bingen, et al. 2009. “Organised Genome Dynamics in the Escherichia Coli Species Results in Highly Diverse Adaptive Paths.” PLoS Genetics 5 (1): e1000344.

Vos, Michiel, and Adam Eyre-Walker. 2017. “Are Pangenomes Adaptive or Not?” Nature Microbiology.

Wickham, Hadley. 2016. ggplot2: Elegant Graphics for Data Analysis. Springer.

Yu, Guangchuang, David K. Smith, Huachen Zhu, Yi Guan, and Tommy Tsan-yuk Lam. 2017. “Ggtree: An R Package for Visualization and Annotation of Phylogenetic Trees with Their Covariates and Other Associated Data.” Edited by Greg McInerny. Methods in Ecology and Evolution / British Ecological Society 8 (1):28–36.

