## Supplementary Figure for "Different evolutionary trends form the twilight zone of the bacterial pan-genome"

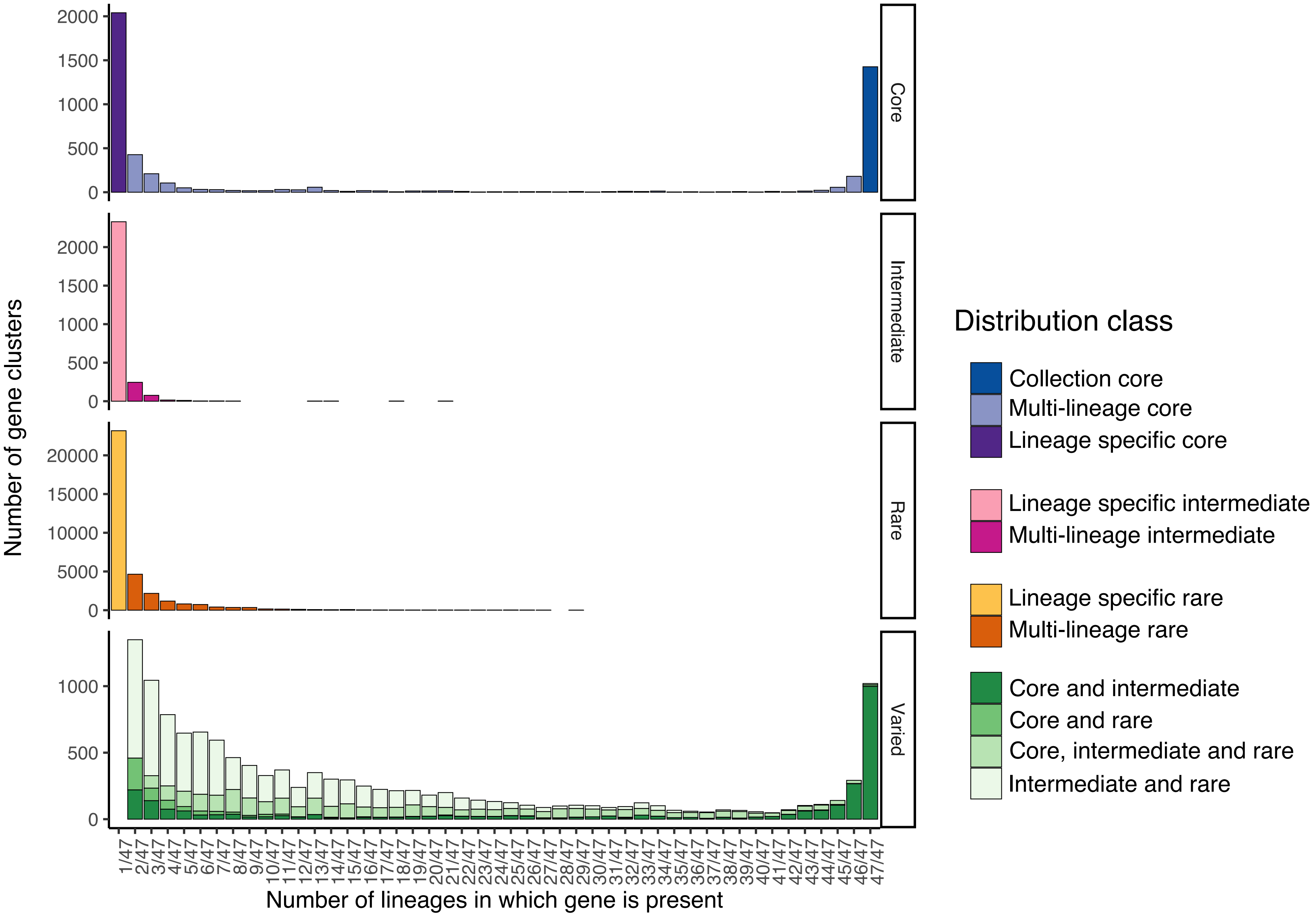


**Figure S1:** Distribution of the number of genes in each distribution class relative to the number of lineages in which they were found.

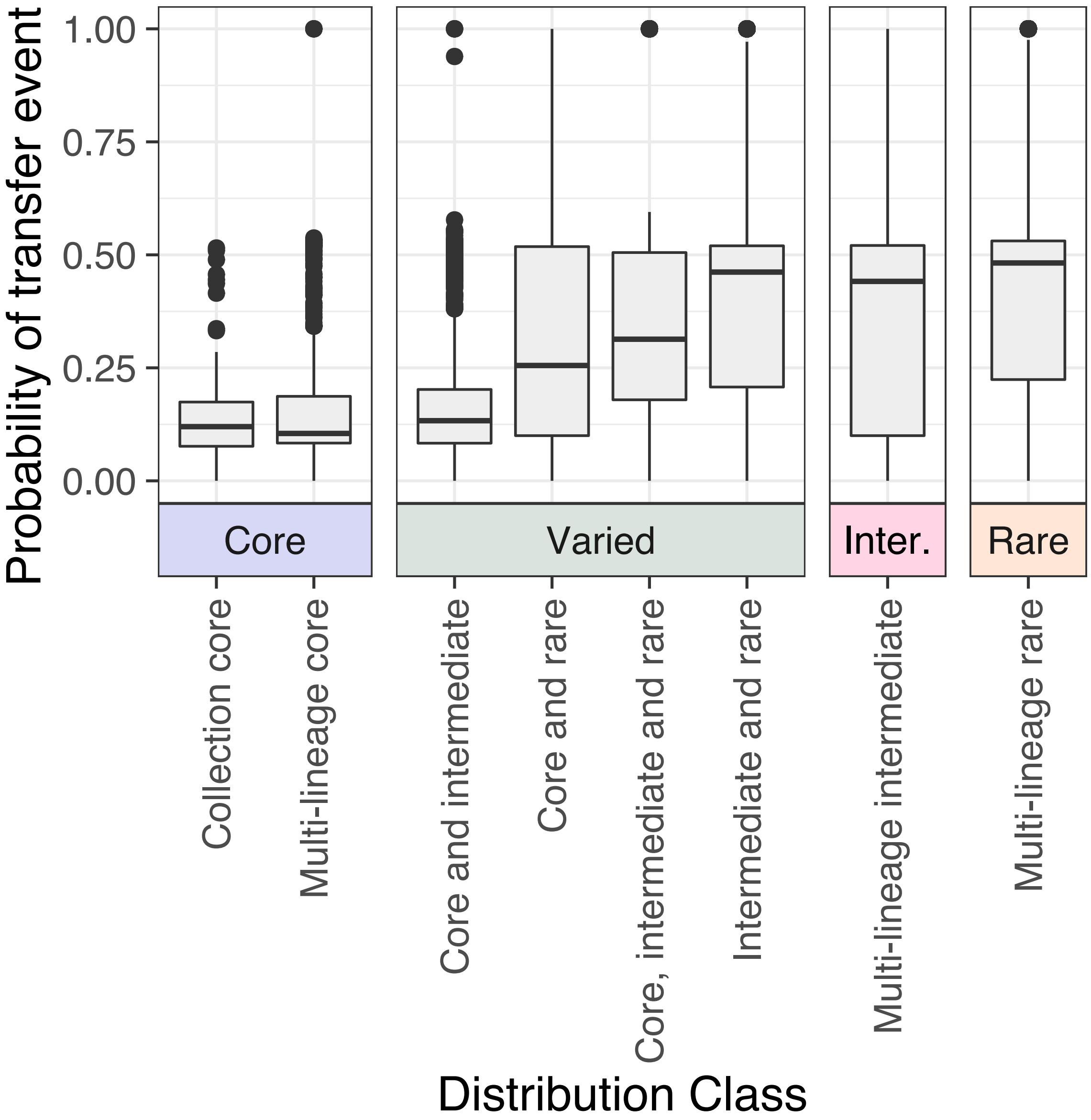


**Figure S2:** Inferred probability of transfer using gene tree/ species tree reconciliation for the genes belonging to the different distribution classes.

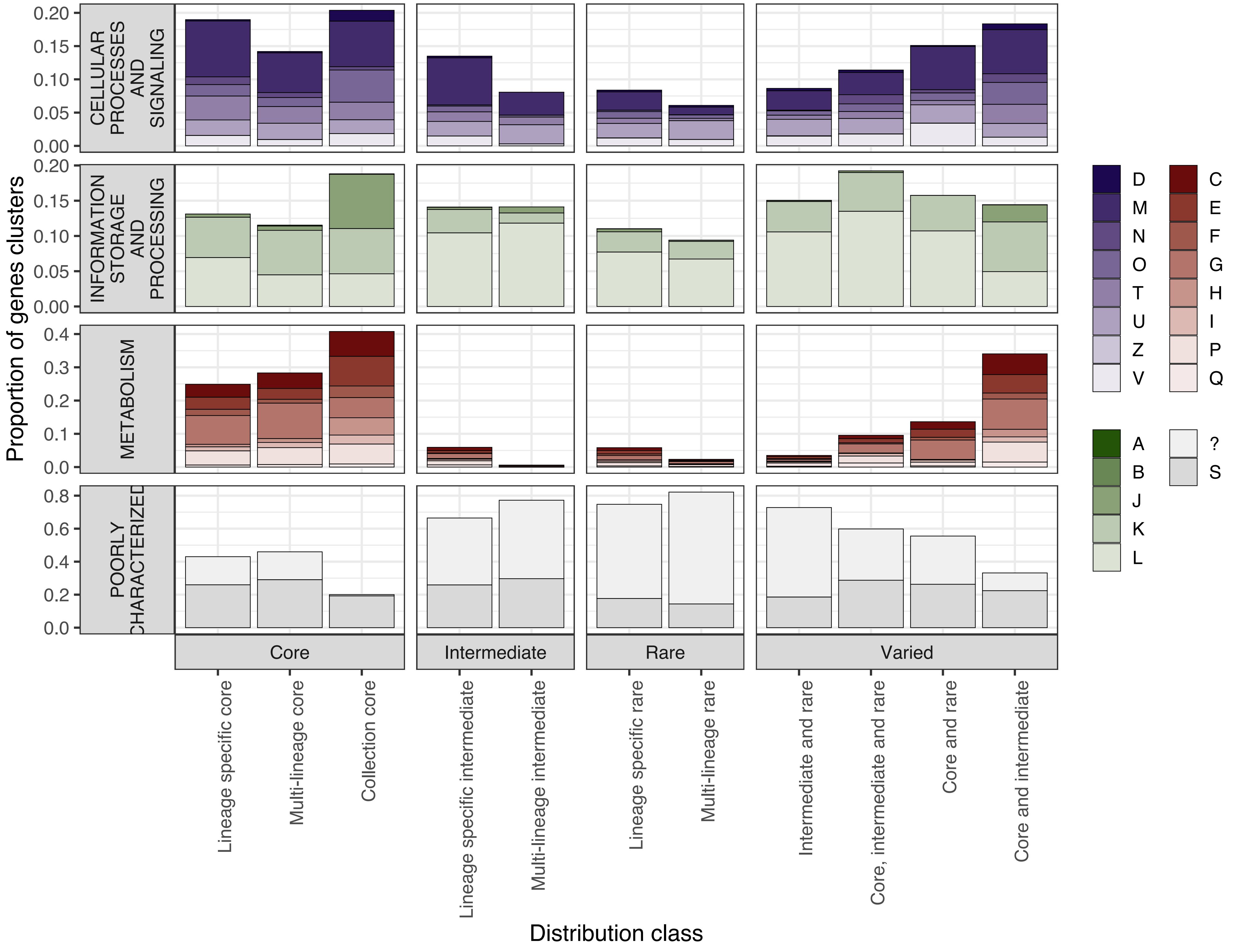


**Figure S3:** Fraction of genes from each occurrence class which were assigned each of the COG categories. D (Cell cycle control, cell division, chromosome partitioning), M ( Cell wall/membrane/envelope biogenesis), N (Cell motility), O (Post-translational modification, protein turnover, and chaperones), T (Signal transduction mechanisms), U (Intracellular trafficking, secretion, and vesicular transport), Z (Cytoskeleton), V (Defence mechanisms), A (RNA processing and modification), B (Chromatin structure and dynamics), J (Translation, ribosomal structure and biogenesis), K (Transcription), L (Replication, recombination and repair), C (Energy production and conversion), E (Amino acid transport and metabolism), F (Nucleotide transport and metabolism), G (Carbohydrate transport and metabolism), H (Coenzyme transport and metabolism), I (Lipid transport and metabolism), P (Inorganic ion transport and metabolism), Q (Secondary metabolites biosynthesis, transport, and catabolism), S (Function unknown) and “?” (unassigned).

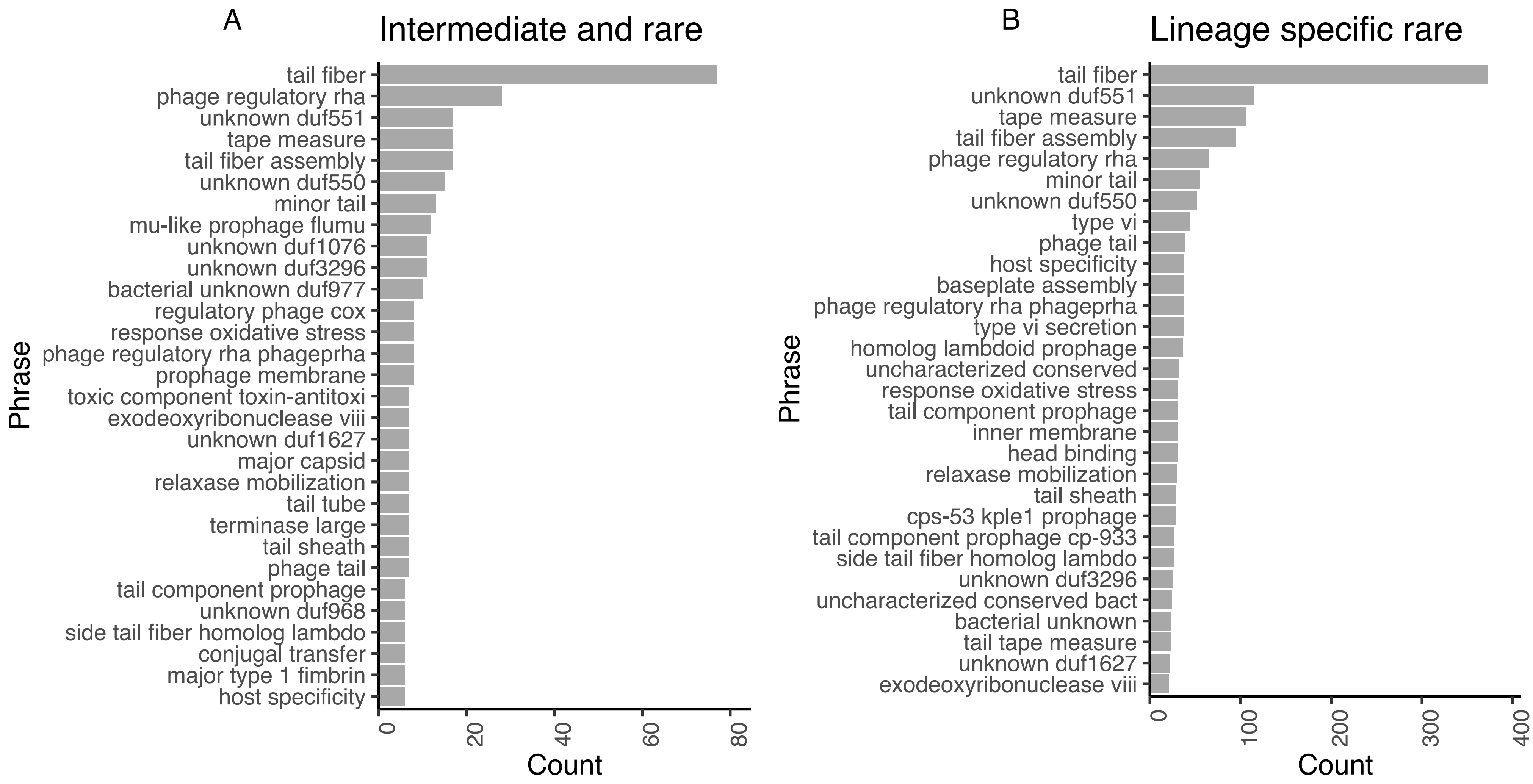


**Figure S4**: Top thirty predicted phrases for genes belonging to the "Poorly Characterised" COG category, for genes belonging to two low frequency distribution classes: 'intermediate and rare' genes (A) and 'lineage-specific rare' genes (B). The majority of the genes in these distribution classes were assigned a COG category of poorly characterised (Supplementary Figure S3).

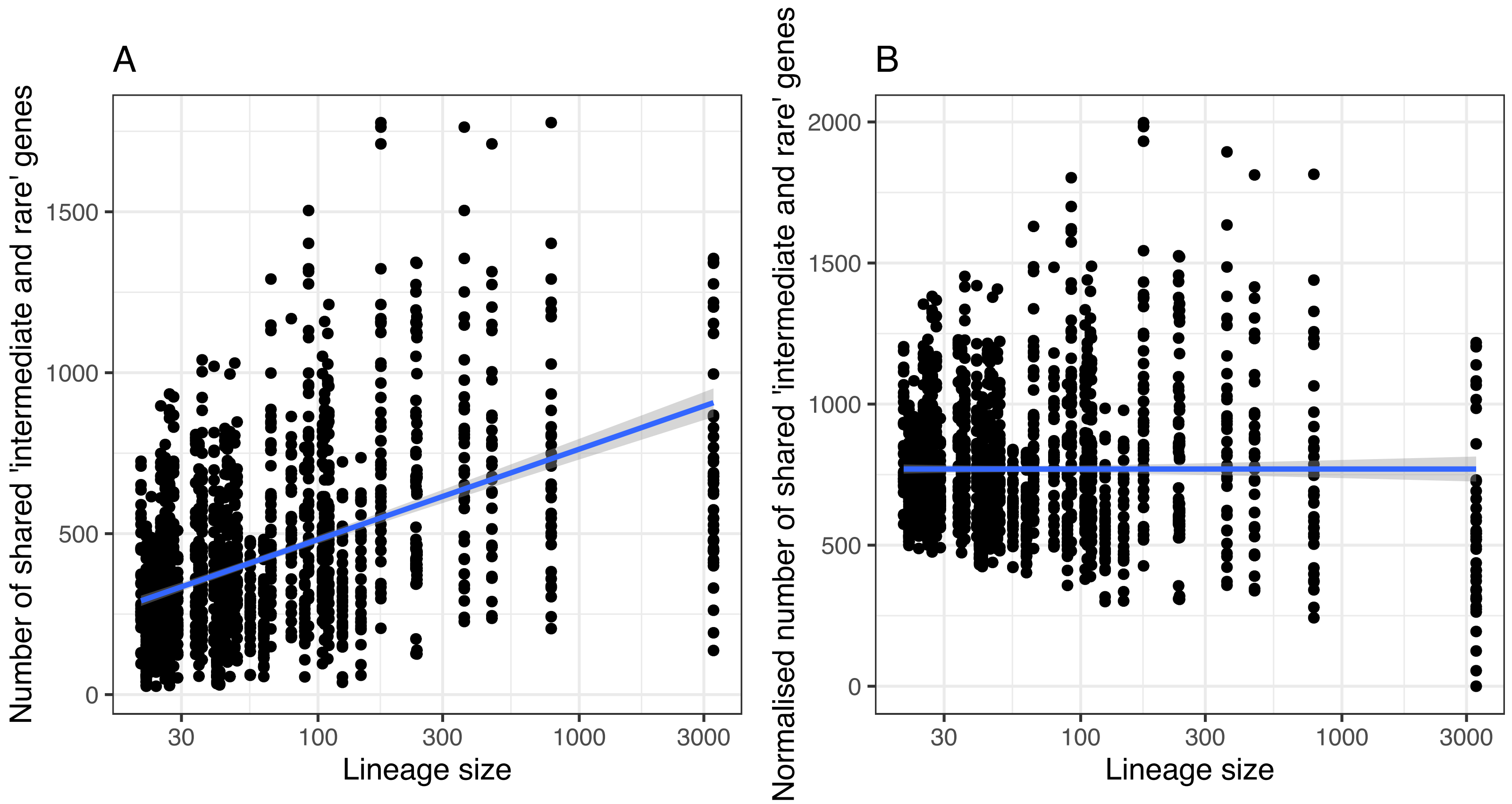


**Figure S5:** Normalisation of shared gene counts to correct for lineage size. A Original value of the number of shared genes between every two lineages, relative to the size of the lineage (log10-transormed). Counts were only considered between lineages belonging to different phylogroups. Line fitted using linear regression, p< 2.2e-16, R2 = 0.22. B Normalised number of shared 'intermediate and rare' genes, corrected based on the fitted line presented in A such that the new counts are

$$counts_{new} = count_{orig} - \beta \times log10(size) - \alpha$$

where beta is the coefficient of the line and alpha is the intercept. Counts were then scaled to be greater than 0. Line presented fitted using linear regression, p=1, R2 = 0.

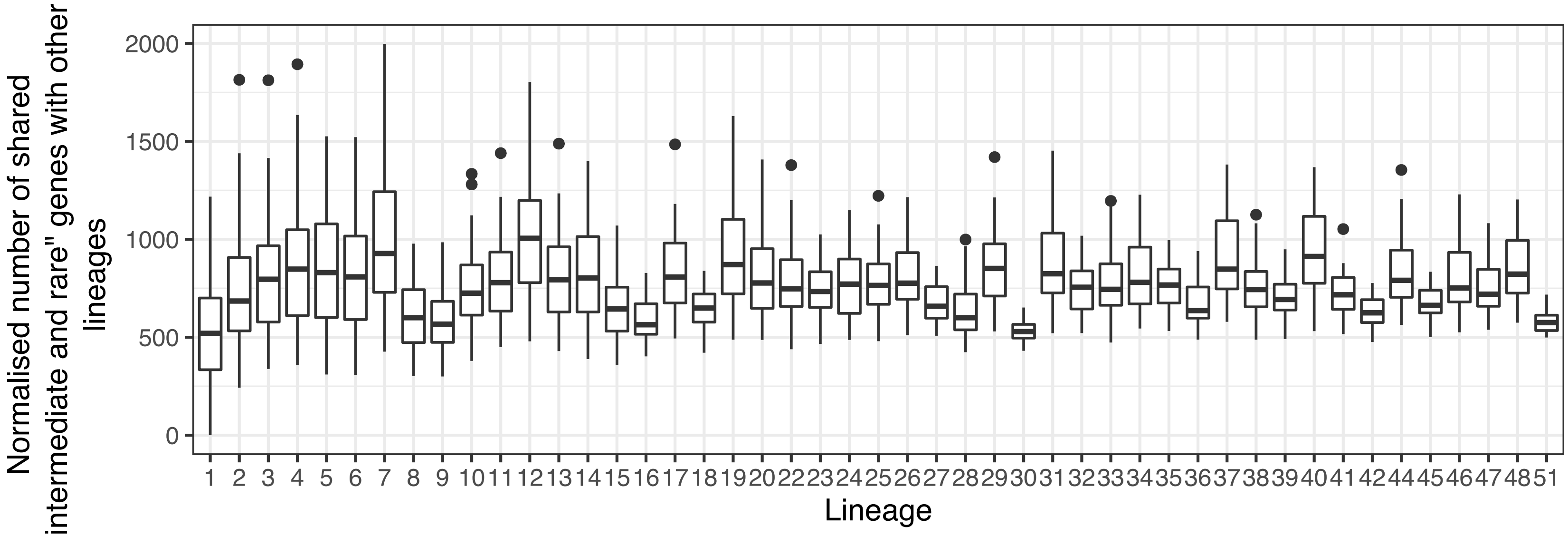


**Figure S6:** Number of shared intermediate and rare genes per isolate, for all lineages. Counts were only considered between lineages belonging to different phylogroups, and were normalised to consider the dependency on the lineage size (Figure S5).

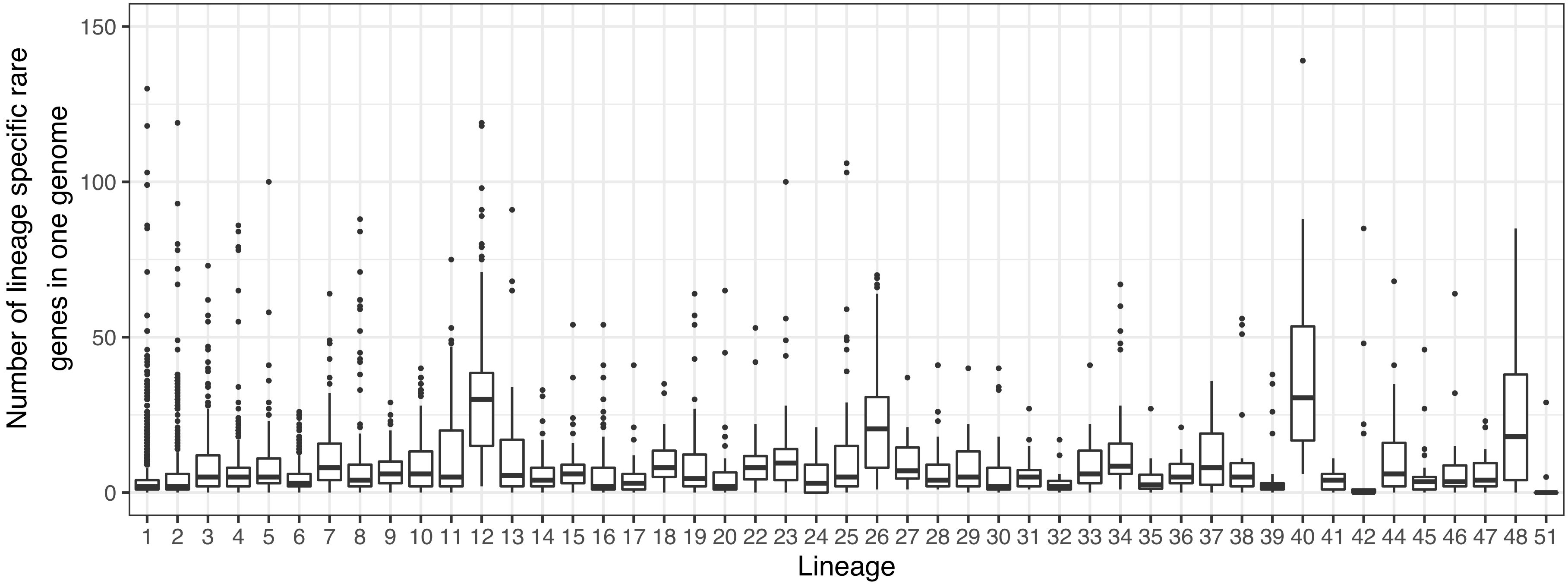


**Figure S7:** Number of ‘lineage specific rare’ genes observed in each isolate, for isolates belonging to each of the lineages
